# Molecular safeguarding of CRISPR gene drive experiments

**DOI:** 10.1101/411876

**Authors:** Jackson Champer, Joan Chung, Yoo Lim Lee, Chen Liu, Emily Yang, Zhaoxin Wen, Andrew G. Clark, Philipp W. Messer

**Author notes:** Corresponding authors (J.C.) (P.W.M).

## Abstract

CRISPR-based gene drives have sparked both enthusiasm and deep concerns due to their potential for genetically altering entire species. This raises the question about our ability to prevent the unintended spread of such drives from the laboratory into a natural population. Here, we experimentally demonstrate the suitability of synthetic target sites and split drives as flexible safeguarding strategies for gene drive experiments.

## INTRODUCTION

In theory, homing gene drives could rapidly spread through a population by copying themselves into a target site on the homologous chromosome, thereby enabling super-Mendelian inheritance^1–7^. Proof-of-principle studies using CRISPR-based drive constructs have now been demonstrated in a variety of potential target systems, ranging from microbes^8^ to insects^9–16^ and mammals^17^.

While some have touted this technology as a potential game-changer in the fight against vector-borne diseases, key questions loom large about our ability to predict the outcome of releasing such a drive into a natural population. Unintended consequences could be catastrophic. At present, these concerns may be hypothetical, given that current drives are inefficient and prone to rapid evolution of resistance ^10,16,18^. However, various approaches for lowering resistance and increasing drive efficiency are already being explored^9,11^. Thus, effective drives may be developed in the near future.

Regardless of the likelihood of their escape and rapid expansion in the field, it is imperative that we safeguard laboratory gene drives so that they cannot accidentally spread into a natural population. Current strategies typically rely on physical confinement of drive-containing organisms. However, it is doubtful whether this is sufficient to completely eliminate the possibility of any accidental escape into the wild. Given that very few escapees can already establish an effective drive in a population^4,19^, additional safety measures should be employed in any experiments with drives potentially capable of spreading.

Two molecular safeguarding strategies have recently been proposed that go beyond physical or ecological confinement^20^. The first strategy involves the targeting of a synthetically-engineered genomic site that is absent in the wild. The second strategy employs a so-called split drive, in which the drive construct lacks its own endonuclease, relying on one engineered into an unlinked site instead. Both strategies should thereby prevent efficient drive outside of their respective laboratory strains. Here, we provide the first experimental demonstration of these two approaches in an insect system and show that they behave similarly to standard drives, suggesting that these strategies can serve as appropriate molecular safeguards in the development and testing of CRISPR gene drives.

## RESULTS & DISCUSSION

### Synthetic target sites

We designed and tested three synthetic target drives in *D. melanogaster*, each targeting an enhanced green fluorescent protein (EGFP) gene introduced at a different genomic site (Figure 1a). To determine conversion efficiencies of these drives (the percentage of EGFP alleles converted to drive alleles in the germline of drive/EGFP heterozygotes), we scored the progeny of crosses between drive/EGFP heterozygotes and EGFP homozygotes. We observed drive conversion efficiencies of approximately 52-54% in females and 32-46% in males (Table 1, SI Dataset S1-S3), which were similar to our previous gene drives^10,11^. We next measured the rate at which ‘r2’ resistance alleles (i.e. those that disrupt the target gene) were formed in the embryo by scoring the progeny of drive/EGFP heterozygous females and EGFP homozygous males. This rate was high in all three drives, ranging from 80-90% (Table 1, Dataset S1). It is thus likely that nearly all EGFP target alleles were converted to resistance alleles. These rates are again similar to what we found for previous drive constructs targeting the autosomal *cinnabar* and X-linked *white* loci^11^, but significantly less than a construct targeting the *yellow*^10^ gene (*p*<0.001, Fisher’s exact test), which we believe is mostly due to location-specific differences in expression levels of Cas9 (and possibly also the gRNA) at the *yellow* locus.

**Table 1.** Drive performances of synthetic target and split drives compared with the standard drives from our previous studies^10,11^.

| Drive | Male drive conversion efficiency | Female drive conversion efficiency | Embryo r2 resistance rate |
| --- | --- | --- | --- |
| EGFP site B | 32±3% | 52±3% | 90±2% |
| EGFP site E | 46±4% | 54±5% | 90±3% |
| EGFP site Y | N/A | 53±3% | 80±2% |
| *cinnabar* | 39±3% | 54±4% | 100±0% |
| *white* | N/A | 59±2% | 77±2% |
| Split-*yellow* | N/A | 74±2% | 74±2% |
| *yellow* | N/A | 63±3% | 20±2% |

**Figure 1.**
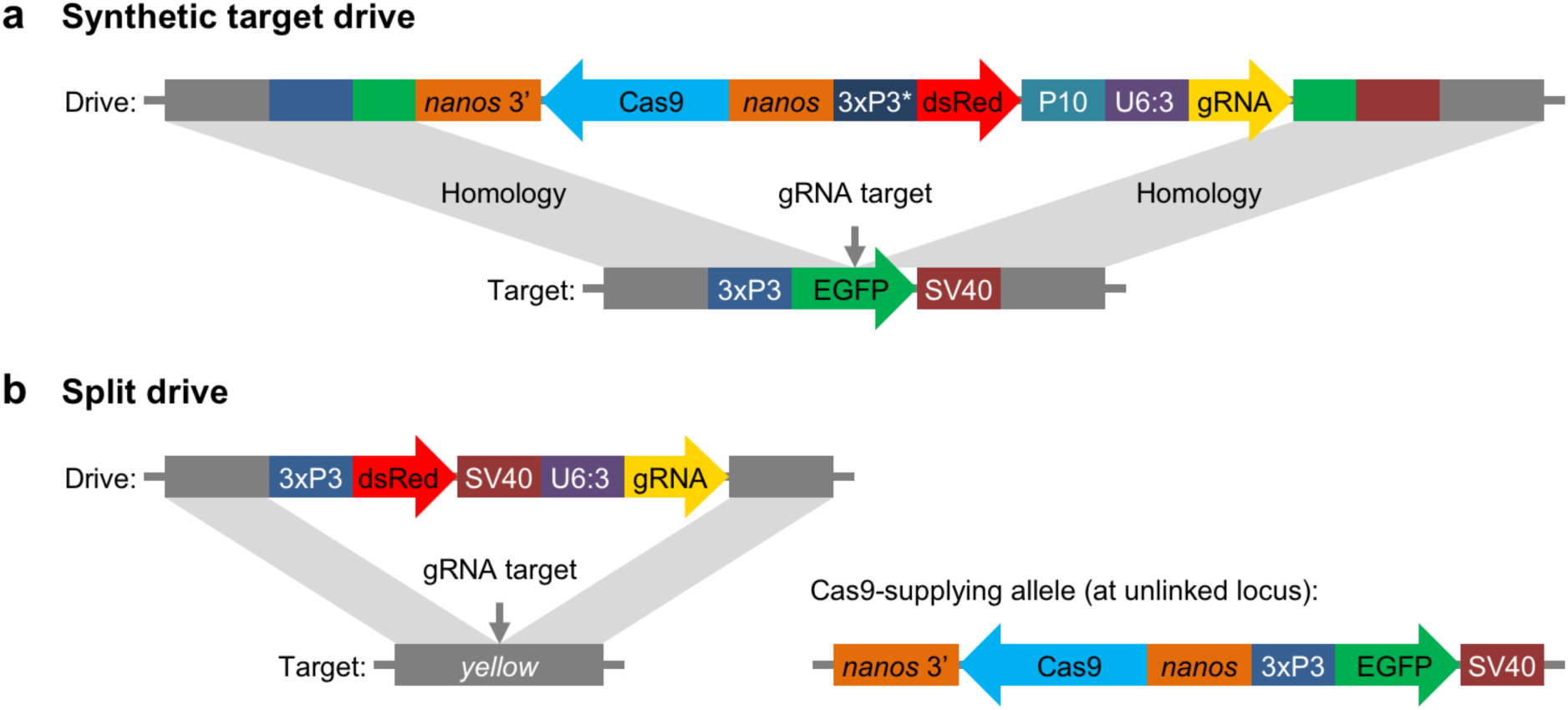
Schematic diagram of our synthetic target drive and split drive constructs. (**a**) The synthetic target drive constructs contain Cas9 with a *nanos* promoter and 3’UTR, a dsRed marker with a slightly recoded (*) 3xP3 promoter and P10 3’UTR, and a gRNA driven by the U6:3 promoter that targets the synthetic EGFP gene. The two homology arms include the EGFP sequence with its 3xP3 promoter and SV40 3’UTR regions. (**b**) The split drive contains a dsRed marker gene driven by a 3xP3 promoter together with a SV40 3’UTR, a gRNA expressed by the U6:3 promoter that targets *yellow*, and two homology arms for *yellow*. The unlinked supporting element contains Cas9 driven by the *nanos* promoter with a *nanos* 3’UTR, and an EGFP marker gene driven by a 3xP3 promoter together with a SV40 3’UTR.

### Split drives

For our split drive system, we designed a drive construct targeting the X-linked *yellow* gene, similar to the one used in a previous study^10^, but lacking Cas9 (Figure 1b). We then designed a second construct containing Cas9 driven by a *nanos* promoter, which was inserted into an attP site on chromosome 2L. We assessed drive performance of this system by crossing males that had the drive element but no Cas9 to females that were homozygous for Cas9 but lacked the drive element. Similarly, we also crossed females homozygous for the drive but lacking Cas9 to males homozygous for Cas9 but lacking the drive. The progeny of these crosses followed Mendelian inheritance rules, indicating that both Cas9 and gRNA must be maternally deposited for resistance alleles to form in the early embryo.

The progeny of *w*^*1118*^ males and drive/wild-type heterozygous females containing one copy of Cas9 were then scored to assess drive conversion efficiency and resistance rates (Table 1, Dataset S4). Compared to our previous results with a standard gene drive targeting *yellow*^10^, where drive conversion efficiency was 62%, we measured a significantly higher drive conversion efficiency of 74% for the split drive (*p*<0.0001, Fisher’s exact test). This improvement may be due to increased efficiency of homology-directed repair for the split drive element compared to the larger standard drive. However, we also observed that early embryo r2 resistance allele formation was much higher in the split drive at 74% compared to the 20% for the standard drive (*p*<0.001, Fisher’s exact test). This is likely because Cas9 at its new site (near synthetic target site B) had higher expression than the Cas9 in the standard drive at *yellow* and that Cas9, rather than the gRNA, is the main limiting factor in determining the resistance rates.

One concern regarding the use of split drives as a surrogate for standard drives is that every genome in the experimental split drive population would contain Cas9, so maternally-deposited Cas9 would likely be present in each embryo, even if the mother did not have a drive element. In combination with the zygotically expressed gRNA from a paternal allele, this might then result in a higher rate of embryo resistance allele formation. However, our finding that both Cas9 and gRNA must be maternally deposited to form such embryo resistance alleles suggests that a split drive in a laboratory population should in fact behave similarly to a standard drive.

A hypothetical split drive where Cas9 is encoded in the driving element and the gRNA forms the supporting element (the reverse of our split drive) would presumably have nearly identical behavior to a standard drive. This is because in this case the Cas9 gene would always be present in the same copy number per individual as in a standard drive, and it would be located at the same genomic position, eliminating the possibility of position-based differences in Cas9 expression levels between the two drives. However, such a strategy would be experimentally less flexible because both elements would have to be redesigned for every new target site, rather than just the drive element when Cas9 is in the supporting element.

The flexibility of the split drive system, facilitated by its genomic separation of Cas9 and gRNA, allowed us to further refine our understanding of the general mechanisms by which homing drives operate. Previous studies have indicated that germline resistance alleles can form in pre-gonial germline cells^11,16^, but it remained unclear whether this could also occur at other stages. The fact that we observed a higher drive conversion efficiency of the split drive compared with a standard drive strongly implies that not all resistance alleles form prior to drive conversion, since resistance allele formation alone should not be affected by the reduced size of the split drive. This raises the possibility that drive conversion could potentially take place as an alternative to resistance allele formation in pre-gonial germline cells, where resistance alleles are known to form. However, a perhaps more likely explanation would be that only a portion of resistance alleles form in pre-gonial germline cells, and that the remainder form either in gametocytes as an alternative to drive conversion or afterward, in late meiosis, when a template for homology-directed repair is no longer available.

We further found evidence that even in individuals that lack any genomic source of Cas9, maternally-deposited Cas9 can persist through to gametocyte formation in the germline, where it can then facilitate successful drive conversion. This was demonstrated by crosses of *w*^*1118*^ males with females that were mosaic yellow and had dsRed from one maternally inherited drive allele. These females received maternally-deposited Cas9 from a heterozygous mother but did not inherit the Cas9 allele themselves, as evidenced by the absence of a EGFP phenotype. Despite lacking a Cas9 gene, these flies nevertheless showed an average drive conversion efficiency of 56% (Dataset S4). By contrast, females from the same cross that were fully yellow, rather than mosaic, and wild-type females showed no detectable drive conversion (in these latter females, ‘r1’ resistance alleles that preserve the function of the target gene had presumably formed at their own embryo stage). It is unlikely that any drive conversion had occurred in these females with full yellow phenotype at the early embryo stage, because in that case their progeny should have displayed biased inheritance of the drive allele. Thus, it appears that when Cas9 and gRNA are maternally-deposited, they can fail to cleave the target site in the early embryo and induce end-joining repair, while nonetheless showing significant cleavage activity in later stages when homology-directed repair is possible. Such Cas9 did not persist to embryos of the subsequent generation, as indicated by the lack of any observed yellow phenotype in female progeny.

We also investigated whether maternally-deposited gRNA was necessary to achieve drive conversion in conjunction with maternally-deposited Cas9. This does not seem to be the case when a genomic source of gRNA is provided. For example, we found that the progeny of drive-heterozygous females without Cas9 (but receiving maternally-deposited Cas9 from a mother heterozygous for Cas9) that had received a paternal drive allele also showed germline drive conversion, with an efficiency of 37% (Dataset S4). An additional 14% of wild-type alleles were converted to r2 resistance alleles, and the remainder most likely remained wild-type. These rates were lower than those of the full split drive, likely because of reduced Cas9 and gRNA activity due to the fact that only maternal and no newly expressed Cas9 could be utilized.

Taken together, our results suggest that in the absence of early embryo resistance alleles, germline drive rates in the next generation may be affected by whether an individual inherits maternally-deposited Cas9 and gRNAs. Individuals receiving a maternal drive allele, as opposed to a paternal allele, will also receive maternally-deposited Cas9, which could increase the level of cleavage during the window for homology-directed repair, thereby increasing the rate of drive conversion. On the other hand, cleavage by maternally-deposited Cas9 prior to this stage in pre-gonial germline cells could form additional resistance alleles compared to individuals with only newly expressed Cas9.

### Conclusions

Our results demonstrate that CRISPR gene drives with synthetic target sites such as EGFP will show highly similar behavior to standard drives and can thus be used for most testing in lieu of these drives. Split drives also show similar performance, while allowing for the targeting of natural sequences in situations where the use of synthetic targets is difficult, such as for certain resistance reduction strategies and population suppression drives that require the targeting of wild-type genes.

We therefore suggest that gene drive research should consistently adopt these molecular safeguarding strategies in the development and testing of new drives. This will be particularly critical for large-scale cage experiments aimed at gaining a better understanding of the expected population dynamics of candidate drives, which will be integral for any informed discussion about their feasibility and risks.

## MATERIALS & METHODS

### Synthetic target drive designs

We constructed three synthetic target drives at different genomic sites (B, E, and Y) into which the EGFP target was inserted. Sites B and E were on chromosomes 2L and 3R, respectively, located 3’ of two protein coding genes. Site Y was on the X chromosome immediately downstream of *yellow*. The synthetic target drives used a slightly recoded 3xP3 promoter (3xP3v2) to drive the dsRed marker and also used a P10 3’UTR. This was to reduce potential misalignment with the 3xP3 promoter and SV40 3’UTR in the homology arms (see Figure 1a), which we found to result in poor drive efficiency in initial tests of drive constructs that used the same 3xP3 promoter and SV40 3’UTR for the EGFP and dsRed markers.

### Genotypes and phenotypes

Since all synthetic target gene drive constructs target inside EGFP, successful insertion of the drives will disrupt this marker. For the split drive, the driving element disrupts *yellow*, causing a recessive yellow body phenotype. If cleavage is repaired by end-joining, rather than homology-directed repair, this will typically result in a mutated target site, creating a resistance allele. Most such resistance alleles will render the target gene nonfunctional due to a frameshift or otherwise sufficient change in the amino acid sequence. We term these alleles ‘r2’. Resistance alleles that preserve the function of the target gene are termed ‘r1’. In some cases, we observed mosaicism for EGFP in the eyes of heterozygotes for the drive and the synthetic target site. This indicates that the *nanos* promoter may drive a low level of expression in somatic cells, though this may be due to its proximity to the 3xP3 promoter for expression in the eyes, since no mosaicism was observed in the body of the split drive flies. The different phenotypes and genotypes of our drive systems are summarized in the SI Dataset, as are calculations for determining drive performance parameters based on phenotype counts.

### Generation of transgenic lines

One line in the study was transformed at GenetiVision by injecting the donor plasmid (ATSabG) into a *w*^*1118*^ *D. melanogaster* line, and seven lines were transformed at Rainbow Transgenic Flies by injecting the donor plasmid (ATSaeG, ATSxyG, BHDgN1bv2, BHDgN1e, BHDgN1y, BHDaaN, IHDyi2) into the same *w*^*1118*^ line. Cas9 from plasmid pHsp70-Cas9gra^21^ (provided by Melissa Harrison & Kate O’Connor-Giles & Jill Wildonger, Addgene plasmid #45945) and gRNA from plasmids BHDaag1, BHDabg1, BHDaeg1, or BHDxyg1 were included in the injection, depending on the target site. For Genetivision injections, concentrations of donor, Cas9, and gRNA plasmids were 102, 88, and 60 ng/μL, respectively in 10 mM Tris-HCl, 23 μM EDTA, pH 8.1 solution. For Rainbow Transgenic Flies injections, concentrations of donor, Cas9, and gRNA plasmids were approximately 350-500, 250-500, and 50-100 ng/μL, respectively in 10 mM Tris-HCl, 100 μM EDTA, pH 8.5 solution. To obtain a homozygous gene drive line, the injected embryos were reared and crossed with *w*^*1118*^ flies. The progeny with dsRed or EGFP fluorescent protein in the eyes, which usually indicated successful insertion of the donor plasmid, were selected and crossed with each other for several generations. The stock was considered homozygous at the drive locus after sequencing confirmed lack of wild-type or resistance alleles.

### Fly rearing and phenotyping

All flies were reared at 25°C with a 14/10 hr day/night cycle. Bloomington Standard medium was provided as food every 2-3 weeks. During phenotyping, flies were anesthetized with CO_2_ and examined with a stereo dissecting microscope. Flies were considered ‘mosaic’ if any discernible mixture of green fluorescence was observed in either eye. However, for flies from synthetic target site drive crosses carrying the drive allele, a fly was only considered mosaic if either eye had less than 50% EGFP phenotype coverage, to avoid identifying flies with possible somatic expression of Cas9 as mosaic for EGFP. This definition was stringent enough that no mosaic insects without the drive were found that would have avoided mosaic classification based on this definition. Fluorescent phenotypes were scored using the NIGHTSEA system only in the eyes (SFA-GR for dsRed and SFA-RB-GO for EGFP). Even though dsRed did bleed through into the EGFP channel, both types of fluorescence could still be easily distinguished.

All experiments involving live gene drive flies were carried out using Arthropod Containment Level 2 protocols at the Sarkaria Arthropod Research Laboratory at Cornell University, a quarantine facility constructed to comply with containment standards developed by USDA APHIS. Additional safety protocols regarding insect handling approved by the Institutional Biosafety Committee at Cornell University were strictly obeyed throughout the study, further minimizing the risk of accidental release of transgenic flies.

### Genotyping

To obtain the DNA sequences of gRNA target sites, individual flies were first frozen and then ground in 30 μL of 10 mM Tris-HCl pH 8, 1mM EDTA, 25 mM NaCl, and 200 μg/mL recombinant proteinase K (Thermo Scientific). The homogenized mixture was incubated at 37°C for 30 min and then 95°C for 5 min. 1 μL of the supernatant was used as the template for PCR to amplify the gRNA target site. DNA was further purified by gel extraction and Sanger sequenced. Sequences were analyzed using the ApE software, available at: http://biologylabs.utah.edu/jorgensen/wayned/ape.

### Plasmid construction

The starting plasmid pCFD3-dU6:3gRNA^22^ (Addgene plasmid #49410) was kindly supplied by Simon Bullock, starting plasmid pJFRC81-10XUAS-IVS-Syn21-GFP-p10^23^ was a gift from Gerald Rubin (Addgene plasmid # 36432), and starting plasmid IHDyi2 was constructed in our previous study^10^. All plasmids were digested with restriction enzymes from New England Biolabs (HF versions, when available). PCR was conducted with Q5 Hot Start DNA Polymerase (New England Biolabs) using DNA oligos and gBlocks from Integrated DNA Technologies. Gibson assembly of plasmids was conducted with Assembly Master Mix (New England Biolabs) and plasmids were transformed into JM109 competent cells (Zymo Research). Plasmids used for injection into eggs were purified with ZymoPure Midiprep kit (Zymo Research). Cas9 gRNA target sequences were identified using CRISPR Optimal Target Finder^24^. Tables of the DNA fragments used for Gibson Assembly of each plasmid, the PCR products with the oligonucleotide primer pair used, and plasmid digests with the restriction enzymes are shown in the Supporting Information.

## ACKNOWLEDGEMENTS

This study was supported by startup funds from the College of Agriculture and Life Sciences at Cornell University to P.W.M, the National Institutes of Health award R21AI130635 to J.C., A.G.C., and P.W.M, and the National Institutes of Health award F32AI138476 to J.C.

